# Capacitation-associated alkalization in human sperm is differentially controlled at the subcellular level

**DOI:** 10.1101/763987

**Authors:** A. Matamoros-Volante, C.L. Treviño

## Abstract

Capacitation in mammalian sperm involves the accurate balance of intracellular pH (pH_i_), but the underlying control mechanisms are not fully understood, particularly regarding the spatiotemporal regulation of the proteins involved in such pH_i_ modulation. Here we employed an image-based flow cytometry technique combined with pharmacological approaches to study pH_i_ dynamics at the subcellular level during sperm capacitation. We found that, upon capacitation induction, sperm cells undergo intracellular alkalization in the head and principal piece regions, but not in the midpiece. The observed localized pH_i_ increases require the initial uptake of HCO_3_^-^, and it is mediated by several proteins acting in a manner consistent with their subcellular localization. Hv1 proton channel and cAMP-activated Protein Kinase (PKA) antagonists impair alkalization mainly in the principal piece. Na^+^/HCO_3_^-^ cotransporter (NBC) and cystic fibrosis transmembrane regulator (CFTR) antagonists impair alkalization only mildly, predominantly in the head. Motility measurements indicate that inhibition of alkalization in the principal piece prevents the development of hyperactivated motility. Altogether, our findings shed light into the complex control mechanisms of pH_i_ and underscore their importance during human sperm capacitation.

**Summary statement:** Human sperm display differential pH_i_ regulation at the subcellular level upon capacitation, involving the participation of PKA kinase signaling pathway and several membrane transport proteins, culminating in hyperactivation.

## INTRODUCTION

The concentration of H^+^ is a ubiquitous regulatory element for most biochemical reactions and it has relevance in many physiological processes, including sperm function (Nishigaki et al., 2014). It has been widely recognized that mammalian sperm must undergo a series of maturation steps in order to develop full fertilizing capabilities; such processes are collectively known as capacitation, and in vivo they only take place once sperm are inside the female reproductive tract (Austin and Sakkas, 1951; Chang, 1951). Intracellular pH (pH_i_) plays a pivotal role in capacitation, controlling various key proteins involved in it. For example, an increase in pH_i_ promotes the activation of KSper currents (Navarro et al., 2007), which are mediated by the SLO3 K^+^ channel (Zeng et al., 2011; Zeng et al., 2013). SLO3 expression is restricted to sperm cells, and *Slo3* knockout male mice are infertile. Although the mechanisms behind this infertility are not completely understood, failure to fertilize is related to a reduction in progressive motility and an impairment of the acrosomal exocytosis process in sperm (Santi et al., 2010).

Additionally, pH_i_ mediates the development of hyperactivated motility, a special kind of sperm movement characterized by asymmetrical flagellar beating, and which is necessary for successful fertilization (Mishra et al., 2018; Suarez, 2008). Such hyperactivation is mediated by CATSPER, a sperm-specific Ca^2+^ channel that is activated by alkaline pH_i_ through interaction with EFCAB9, a pH-tuned Ca^2+^ sensor that controls CATSPER gating (Hwang et al., 2019). Notably, loss-of-function mutations on any of the CATSPER channel subunits cause infertility in male mice (Qi et al., 2007; Ren et al., 2001) and humans (Avenarius et al., 2009), mainly due to the inability of sperm to hyperactivate. It is widely recognized that pH_i_ regulation in mice sperm involves the participation of another sperm-specific protein, the Na^+^/H^+^ exchanger (sNHE), which drives H^+^ extrusion employing the cell’s [Na^+^] gradient. Similar to CATSPER and SLO3, the lack of sNHE results in male infertility (Wang et al., 2003). While humans express an orthologous sNHE gene, its involvement in human sperm physiology remains elusive. In this regard, the proton channel (Hv1) has been proposed as the main pH_i_ regulator during human sperm capacitation and hyperactivation (Lishko et al., 2010), and its activity has been linked to the activation of CATSPER, leading to Ca^2+^ influx and the concomitant changes in motility patterns (Lishko and Kirichok, 2010; Miller et al., 2015) (Lishko and Kirichok, 2010; Miller et al., 2018). Interestingly, the subcellular localization of all aforementioned proteins is restricted to the flagellum, particularly the principal piece region, which is consistent with their role in motility. On the other hand, our group and others have described the expression and participation of an additional set of proteins during mammalian sperm capacitation, which are related to HCO_3_^-^ transport and thus could potentially participate in pH_i_ balance as well. These proteins include members of the SLC26 (Chávez et al., 2012; El Khouri et al., 2018) and SLC4 (Demarco et al., 2003; Parkkila et al., 1993; Puga Molina et al., 2018; Zeng et al., 1996) HCO_3_^-^ transporter families. One member of the SLC4 family, namely the electrogenic Na^+^/HCO_3_^-^ cotransporter (NBC), appears to mediate HCO_3_^-^ influx, which is required for downstream activation of signaling networks essential for capacitation, such as the cAMP-activated Protein Kinase (PKA) pathway (Demarco et al., 2003; Puga Molina et al., 2018). This particular pathway also seems to mediate plasma membrane hyperpolarization, a hallmark of capacitation, via stimulation of the Cystic Fibrosis Transmembrane Regulator (CFTR) Cl^-^/HCO_3_^-^ channel (Chávez et al., 2012; Hernández-González et al., 2007; Puga Molina et al., 2017). Interestingly, pharmacological blocking of CFTR impairs capacitation in mice (Li et al., 2010; Xu et al., 2007) and human (Puga Molina et al., 2017) sperm. Also, its genetic ablation produces subfertility in mice (Xu et al., 2007). Notably, the subcellular localization of these HCO_3_^-^ transporters differs from that of the H^+^ extruders (i.e. Hv1 and sNHE), with some of the former being mainly localized in the head, and to some extent in the midpiece, but not in the principal piece (Liu et al., 2012; Nishigaki et al., 2014). This suggests that pH_i_ might be differentially regulated throughout the cell, presumably through the participation of different proteins.

A few studies have provided evidence of the net pH_i_ increase that occurs during capacitation, by measuring initial and end point pH_i_ (Cross and Razy-Faulkner, 1997; Lopez-Gonzalez et al., 2014). But despite the importance of pH_i_ in sperm physiology, there have been no examinations of pH_i_ kinetics throughout the entire capacitation process, nor have they been tracked in distinct sperm cell regions. Conducting such studies in sperm cells poses unique experimental challenges given their complex morphology, motility and asymmetrical anatomy, which results in highly compartmentalized physiological cell signals (Buffone et al., 2012).

We recently developed a completely novel strategy to analyze intracellular events in a statistically relevant number of cells, using image-based flow cytometry along with a segmentation process that provides spatial resolution within individual sperm cells (Matamoros-Volante et al., 2018). In the present work, we employed this technique to investigate pH_i_ kinetics at the subcellular level during human sperm capacitation. We found that, upon the cells’ contact with capacitation medium, pH_i_ remained constant in the midpiece, while it increased in the head and in the principal piece, displaying different kinetics. Using pharmacology, we found that multiple proteins mediate the observed pH_i_ changes, with their involvement being distinct in the head and in the principal piece. Lastly, motility measurements indicated that these proteins are required for hyperactivation, but not to maintain total motility. Altogether, our results suggest that pH_i_ modulation in human sperm involves the participation of an entire set of proteins, with the pH_i_ changes being orchestrated in a localized, and possibly time-dependent fashion.

## RESULTS

### During capacitation, pH_i_ increases in the head and in the principal piece, but not in the midpiece

While a pH_i_ increase has been widely recognized as a hallmark of sperm capacitation (Nishigaki et al., 2014), the dynamics and subcellular localization of this alkalization, to our knowledge, had not been previously explored. We employed the pH-sensitive fluorescent probe BCECF to track subcellular pH_i_ changes in human sperm cells using an image-based flow cytometer (Fig. S3A-B). To demonstrate that our previously reported segmentation process (Matamoros-Volante et al., 2018) was suitable for measuring pH_i_ changes in distinct sperm cell regions (Fig. S3C-D), we exposed BCECF-loaded sperm cells to an alkalizing agent (20 mM trimethylammonium, TMA) known to produce a sustained pH_i_ increase of around 0.4 units (Alasmari et al., 2013). As described in Materials and Methods, cell regions were arbitrarily considered to have a high pH_i_ if their fluorescence value was higher than those of the third quartile in the NC condition. Then, in order to determine whether any given treatment had an alkalizing effect on pH_i_, the change in the percentage of subcellular regions with high pH_i_ (Δ%) was calculated with respect to that of the corresponding NC condition, after having eliminated outliers (i.e. those in the top 5%). When cells were exposed to 20 mM TMA, we observed a reproducible increase in the percentage of cells exhibiting high pH_i_ (Δ%) in each of the three distinct subcellular regions analyzed, i.e. head, midpiece and principal piece (Fig. S3E). The pooled fluorescence values for all subcellular regions analyzed for all sperm donors are also shown as boxplots (Fig. S3F). With both approaches used for data analysis, the observed localized pH_i_ increases caused by TMA exposure were statistically significant for all three cell regions, confirming the reliability of our technique

We then studied the pH_i_ dynamics in each subcellular region during capacitation, triggered by exposure to capacitation medium (Fig. 1). Representative images of cells at different capacitation time points are shown in Fig. 1A. As seen in Fig. 1B, there was a significant increase in the percentage of cells with high head pH_i_ (hd-pH_i_) after the initial exposure to capacitation medium (t=0 min), (Δ%=17, *p*<0.0001). Δ% reached a maximum after 15 minutes (Δ%=21, *p*<0.0001), and although it gradually dropped and leveled off at Δ%∼16, the increase in the percentage of cells with high hd-pH_i_ was statistically significant up to 240 minutes of capacitation (*p*<0.0001). Similar dynamics were observed when the pH_i_ was measured in the principal piece (pp-pH_i_), though the changes in Δ% were not as pronounced (Fig. 1B). The increase in the percentage of cells with high pp-pH_i_ became statistically significant only after 15 minutes of capacitation (Δ%=10, *p*=0.0023), reaching a maximum at 30 minutes (Δ%=15, *p*=0.003). Δ% value then dropped to ∼9 after 45 minutes and remained essentially constant up to 240 minutes of capacitation, though the increase in Δ% was no longer statistically significant throughout this time period. In contrast, the change in the percentage of cells with high midpiece pH_i_ (mp-pH_i_) consisted overall of a slight and gradual decrease throughout the entire capacitation period analyzed, though the observed differences were never statistically significant (Fig. 1B). These results display similar statistics when the pooled fluorescence values for all the subcellular regions measured from all donors are analyzed as boxplots (Fig. 1C). The fact that, unlike the other two subcellular regions, the mp-pH_i_ remained unchanged was rather surprising to us, and even though we were able to detect a statistically significant pH_i_ increase in the midpiece using TMA (Fig. S3E), we wanted to verify that this was not simply due to a limitation in our experimental methodology, which could potentially be preventing the reliable detection of fluorescence changes in the midpiece. To this end, we incubated sperm cells with 250 nM MitoTracker Green FM, a mitochondrial-specific fluorescent dye that has been employed as a marker for membrane mitochondrial potential (Sousa et al., 2011). We then triggered a change in fluorescence by challenging these cells with 1 μM CCCP (carbonyl cyanide m-chlorophenyl hydrazine), a mitochondrial electron transport system disrupter. A clear reduction of MitoTracker fluorescence was detected in the midpiece (data not shown), thus indicating that our mp-pH_i_ measurements are reliable. Given that no significant changes in mp-pH_i_ were detected during capacitation, we decided to analyze only the head and principal piece regions in all further pH_i_ dynamics studies.

**Figure 1.**
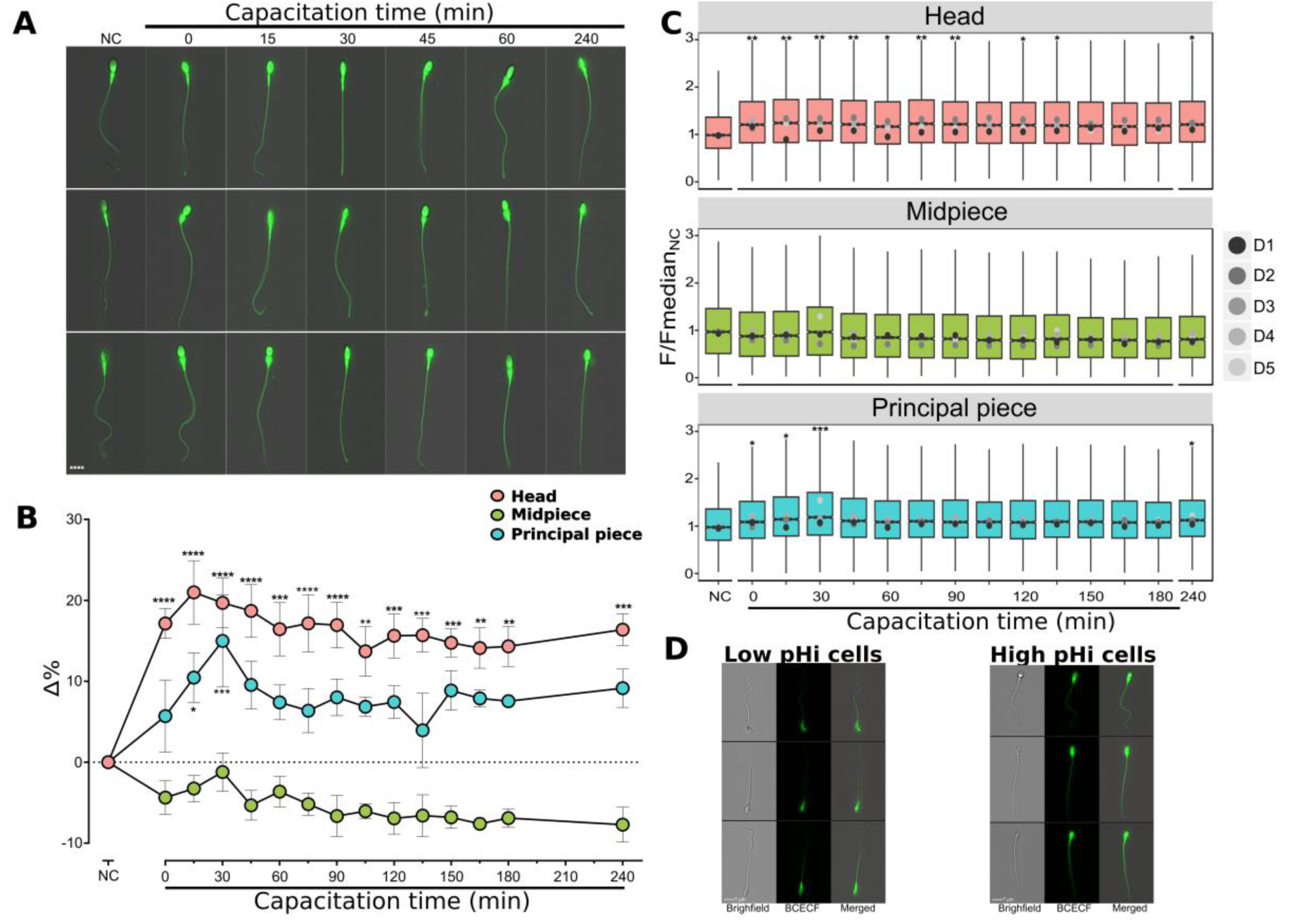
Intracellular alkalization during sperm capacitation occurs in the head and the principal piece but not in the midpiece. **A.** Three representative fluorescence images of BCECF-stained human sperm cells either non-capacitated (NC) or at the indicated capacitation times. **B.** Δ% values (Δ%=%T-%NC) for each subcellular region in cells either non-capacitated (NC) or at the indicated capacitation times. Data represent the mean ± s.e.m. (n=5). **C.** Boxplots of normalized BCECF fluorescence values for each subcellular region in cells either non-capacitated (NC) or at the indicated capacitation times. Gray-scale dots (D1-D5) within boxplots indicate the median fluorescence value for the entire sperm population from each donor, consisting of at least 1,000 cells each (n=5). Data were compared by one-way ANOVA considering capacitation time as one factor. Tukey’s multiple comparison test was employed as post hoc analysis. \**p*<0.05, \*\**p*<0.01, \*\*\**p*<0.001, \*\*\*\**p*<0.0001. **D.** Three representative images of cells considered as having high or low pH_i_. Figure 2

Before proceeding any further, however, we wanted to verify that our experimental conditions were not causing cell damage due to BCECF phototoxicity, even though this was not expected to occur since cells were illuminated for a very short time (milliseconds) during data acquisition. However, BCECF was present in the cell samples up to 6 hours of capacitation, with aliquots being taken for measurements at different time points. To explore whether this exposure was deleterious to the cells, we used PI as a marker for viability. We did not find change in the percentage of viable cells after either 1 or 6 hours of incubation with BCECF, compared to the NC unstained cells (Fig. S4A). Additionally, we wanted to exclude the possibility that any measured pH_i_ increases were simply caused by the time that cells spent in incubation. To this end, we incubated cell samples during 1 and 6 hours in a NC medium. As seen in Fig. S4B-C, no significant changes in pH_i_ were observed at either of these two time points in any of the three subcellular regions. Altogether, these data indicate that the observed pH_i_ increases are a result of incubation in the presence of capacitation medium.

### Absence of HCO_3_^-^ or blockage of HCO_3_^-^ influx prevent pH_i_ increases during capacitation in both the head and the principal piece

When sperm are ejaculated they are exposed to a higher extracellular concentration of HCO_3_^-^ (Owen and Katz, 2005), which is mimicked *in vitro* through exposure to capacitation medium (25 mM HCO_3_^-^, similar to the concentration found in the seminal fluid and the female reproductive tract). To explore the role of HCO_3_^-^ in the observed pH_i_ changes, we incubated sperm in either an incomplete capacitation medium lacking HCO_3_^-^ (no HCO_3_^-^), or in complete capacitation medium containing 100 µM DIDS, a general inhibitor of anionic transporters, to block HCO_3_^-^ entry through channels and transporters at three capacitation times (0, 60 and 240 min). Under both conditions, the pH_i_ increase was completely abolished in the head and in the principal piece (Fig. 2A-C, Fig. S2A). Statistical analysis of both the average Δ%A values and the pooled fluorescence values for all subcellular regions analyzed from all donors yielded comparable statistically significant differences. These results suggest that HCO_3_^-^ uptake via anionic transporters is necessary to induce the rise in pH_i_ in both sperm regions.

**Figure 2.**
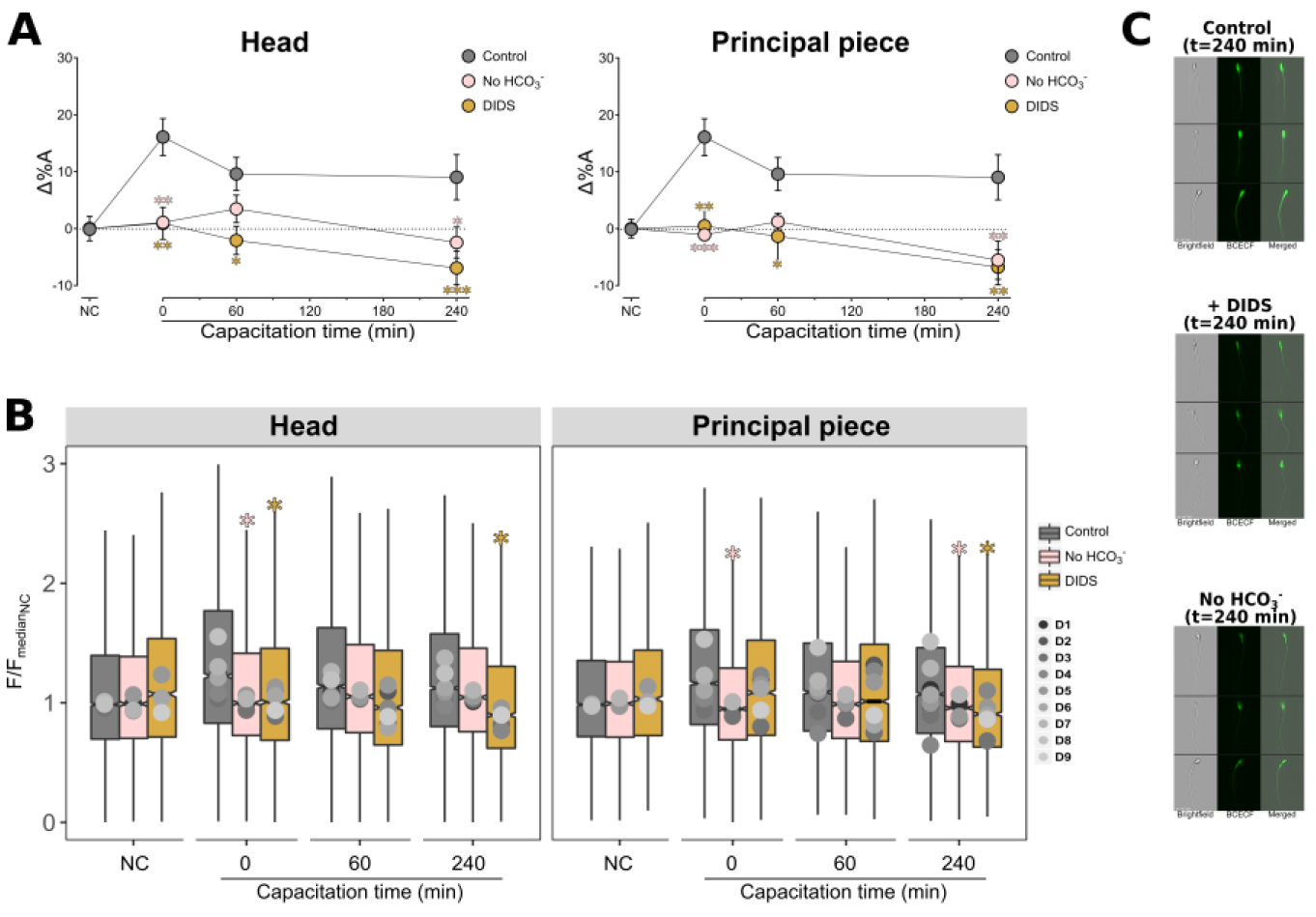
HCO_3_^-^ influx is required for the initial and sustained pH_i_ increase in the head and the principal piece. **A.** Δ%A values (Δ%A=%T-%NC_A_) for each subcellular region from human sperm cells placed in non-capacitating medium or in complete capacitating medium and in both media in the presence or absence of 100 µM DIDS, or medium lacking HCO_3_^-^ (applies only to capacitating medium). Values were measured at the indicated capacitation times. Data represent the mean ± s.e.m. (n=9 for Control, n=6 for DIDS, n=4 No HCO_3_^-^). **B.** Boxplots of normalized BCECF fluorescence values for each subcellular region in cells under the same conditions as in **A**. Gray-scale dots (D1-D9) within boxplots indicate the median fluorescence value for the entire sperm population from each donor, consisting of at least 1,000 cells each (n=9 for Control, n=6 for DIDS, n=4 No HCO_3_^-^). **C**. Three representative images of cells capacitated for 240 min under the indicated conditions. Data were compared using two-way ANOVA with incubation time as one factor, and treatment as the other factor. \**p*<0.05, \*\**p*<0.01, \*\*\**p*<0.001.

### NBC and CFTR have a minor role in cytoplasmic alkalization during capacitation in the head, but not in the principal piece

Previous results from our group have demonstrated that upon HCO_3_^-^ exposure, sperm cells become hyperpolarized due to HCO_3_^-^ uptake mediated by an electrogenic NBC (Demarco et al., 2003; Puga Molina et al., 2018). We thus explored whether pharmacological inhibition of NBC proteins could also prevent pH_i_ increases during capacitation. Interestingly, the mere pre-incubation (10 min) of sperm under NC conditions with a specific antagonist of NBC proteins (S0859, 5 µM) (Ch’en et al., 2008), provoked acidification in the head (Δ% = −11, *p* = 0.0096) compared to NC conditions (Fig. S2B). During capacitation, NBC blockage did not prevent the pH_i_ increases in the head nor in the principal piece (Fig. 3A-B).

**Figure 3.**
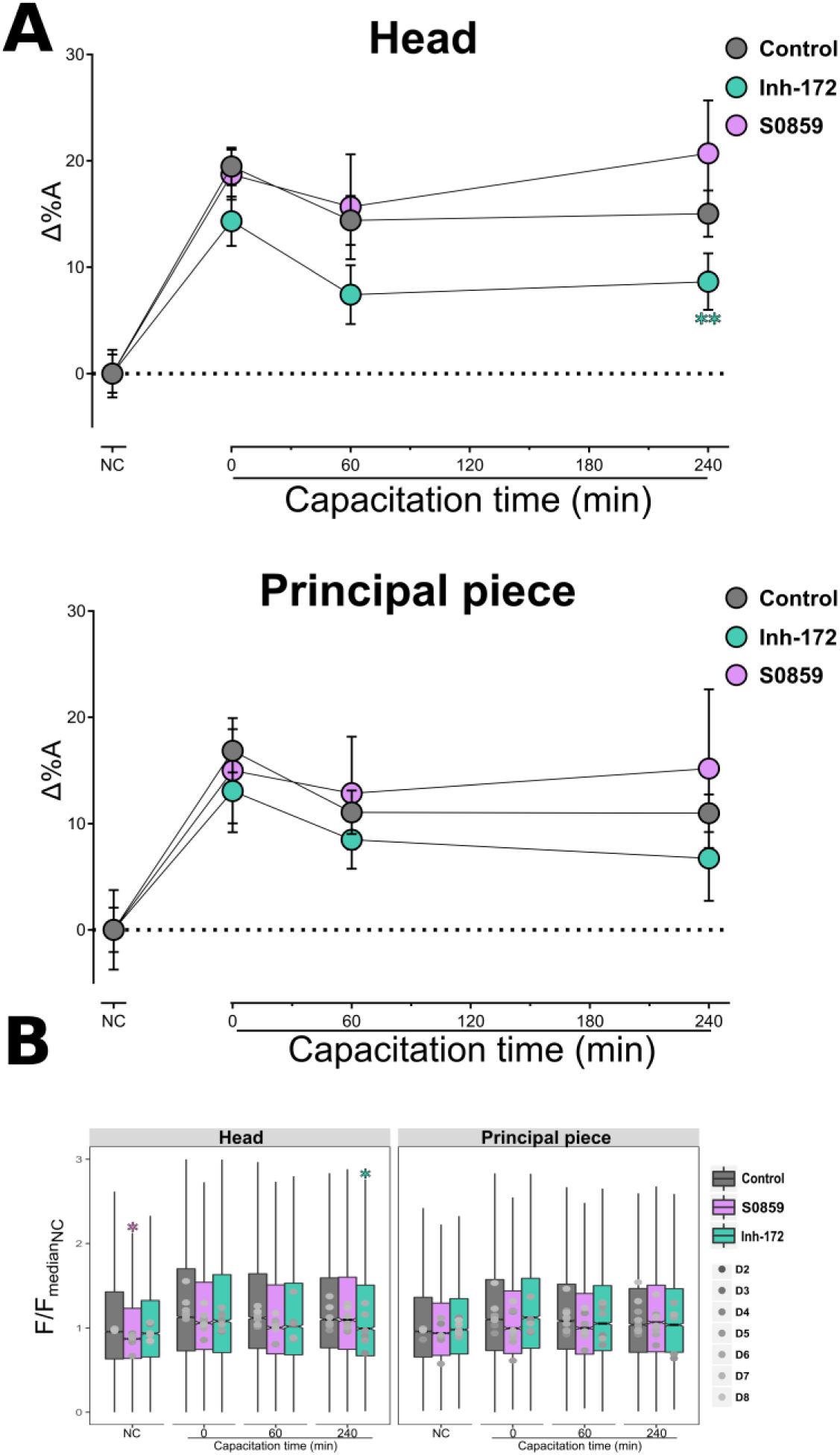
NBC and CFTR have a minor role in cytoplasmic alkalization during capacitation. **A.** Δ%A values (Δ%A=%T-%NC_A_) for each subcellular region from human sperm cells placed in non-capacitating medium or in complete capacitating medium and in both media in the presence or absence of 5 µM S0859 or Inh172. Values were measured at the indicated capacitation times. Data represent the mean ± s.e.m. (n=8 for Control and Inh-172, n=6 for S0859). **B.** Boxplots of normalized BCECF fluorescence values for each subcellular region in cells under the same conditions as in **A**. Gray-scale dots (D1-D8) within boxplots indicate the median fluorescence value for the entire sperm population from each donor, consisting of at least 1,000 cells each (n=8 for Control and Inh-172, n=6 for S0859). Data were compared using two-way ANOVA with incubation time as one factor, and treatment as the other factor. \**p*<0.05, \*\**p*<0.01.

Previously, we reported that pharmacological blocking of CFTR produces cytoplasmic acidification after 5 hours of capacitation (Puga Molina et al., 2017). Since this channel can also transport HCO_3_^-^ into the cell, we used Inh-172, a specific CFTR antagonist, to explore the role of CFTR in the observed alkalization of subcellular regions. During capacitation, there is a decrease in the percentage of cells with high hd-pH_i_, though it is statistically significant only at 240 min (Fig. 3A-B). Inhibition of CFTR did not affect the pH_i_ increase in the principal piece (Fig. 3A-B, Fig.S2B). Although pre-incubation with Inh-172 also caused a reduction in Δ% under NC conditions both in the head and principal piece, it was not statistically significant in these cases (Fig. S2B)

### PKA signaling pathway participates in the regulation of capacitation-associated alkalization in the principal piece, but not in the head

The above observations suggest that HCO_3_^-^ influx is indispensable for the initial and sustained pH_i_ increases, which are stable during at least during 4 hours of capacitation. HCO_3_^-^ is a key component of capacitation medium, and is known to activate a PKA pathway leading to important changes during capacitation (Buffone et al., 2014). It is well accepted that a HCO_3_^-^ influx stimulates cAMP production via a HCO_3_^-^-sensitive adenylyl cyclase (ADCY10) with the subsequent PKA activation (Okamura et al., 1985). Additionally, Puga-Molina, et al. (2017), showed that pharmacological blocking of PKA with H89 induces cytoplasmic acidification in human sperm, measured after 5 h of capacitation. We wondered whether the HCO_3_^-^ requirement for alkalization that we observed is linked to PKA pathway activation. We tested this possibility by incubating sperm with either H89 (30 µM), a PKA inhibitor, or KH7 (50 µM) an ADCY10 antagonist. While neither inhibitor affected Δ%A in the head during capacitation (Fig. 4A-B), both diminished it in the principal piece, though the decrease was statistically significant only at 240 min (*p*=0.032 and 0.0285 respectively) (Fig. 4A). However, when fluorescence data are compared through boxplots (Fig. 4B), the reduction in the BCECF fluorescence in the principal piece was statistically significant at all capacitation time points (*p*<0.0306) for H89, and only at 240 min for KH7 (*p*=0.0182). Additionally, preincubation under NC conditions with H89 reduced Δ% of cells with high pp-pH_i_ (Fig. S2C).

**Figure 4.**
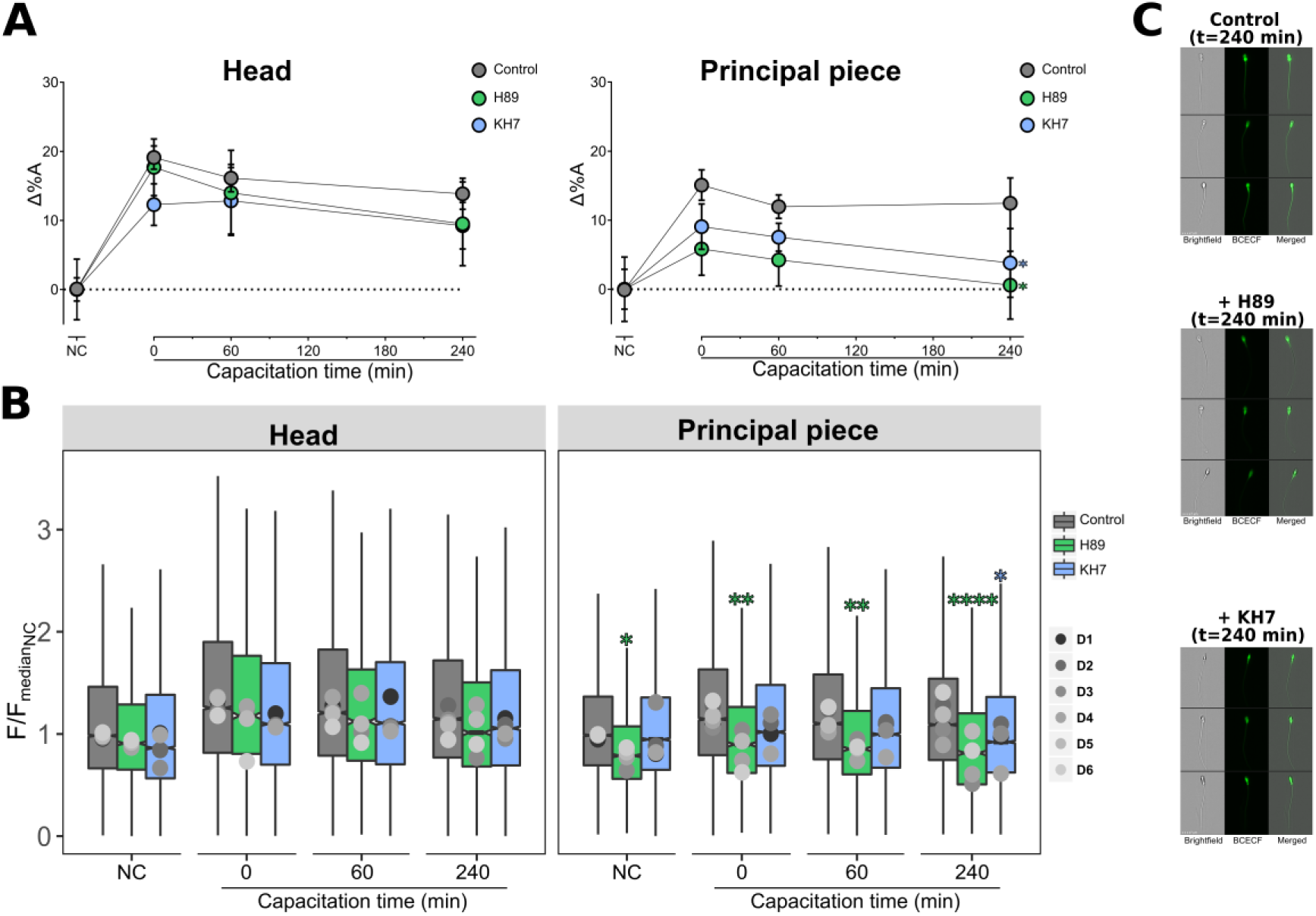
The PKA signaling pathway participates in the regulation of capacitation-associated alkalization in the principal piece, but not in the head. **A.** Δ%A values (Δ%A=%T-%NC_A_) for each subcellular region from human sperm cells placed in non-capacitating medium or in complete capacitating medium and in both media in the presence or absence of 50 µM KH7 or 30 µM H89. Values were measured at the indicated capacitation times. Data represent the mean ± s.e.m. (n=6 for Control, n=4 for KH7 and H89). **B.** Boxplots of normalized BCECF fluorescence values for each subcellular region in cells under the same conditions as in **A**. Gray-scale dots (D1-D6) within boxplots indicate the median fluorescence value for the entire sperm population from each donor, consisting of at least 1,000 cells each (n=6 for Control, n=4 for KH7 and H89). Data were compared using two-way ANOVA with incubation time as one factor, and treatment as the other factor. \**p*<0.05, \*\**p*<0.01, \*\*\*\**p*<0.0001.

### Inhibition of Hv1 prevented alkalization in the principal piece but not in the head

Previous reports have demonstrated that, upon capacitation, Hv1 activity increases in human but not in mice sperm (Lishko et al., 2010). We thus tested whether Hv1 were involved in the observed increases in hd-pH_i_ and pp-pH_i_ by incubating cells with 200 µM Cl-GBI, a specific Hv1 antagonist (Hong et al., 2014). Cl-GBI had no significant effect on Δ%A in the head (Fig. 5A-B, Fig. S2D). Cl-GBI, induced a strong reduction in Δ%A in the principal piece at all incubation times (*p*<0.0387) (Fig. 5A-B) as well as an acidification when preincubated under NC conditions (*p*=0.0482) (Fig. S2D).

**Figure 5.**
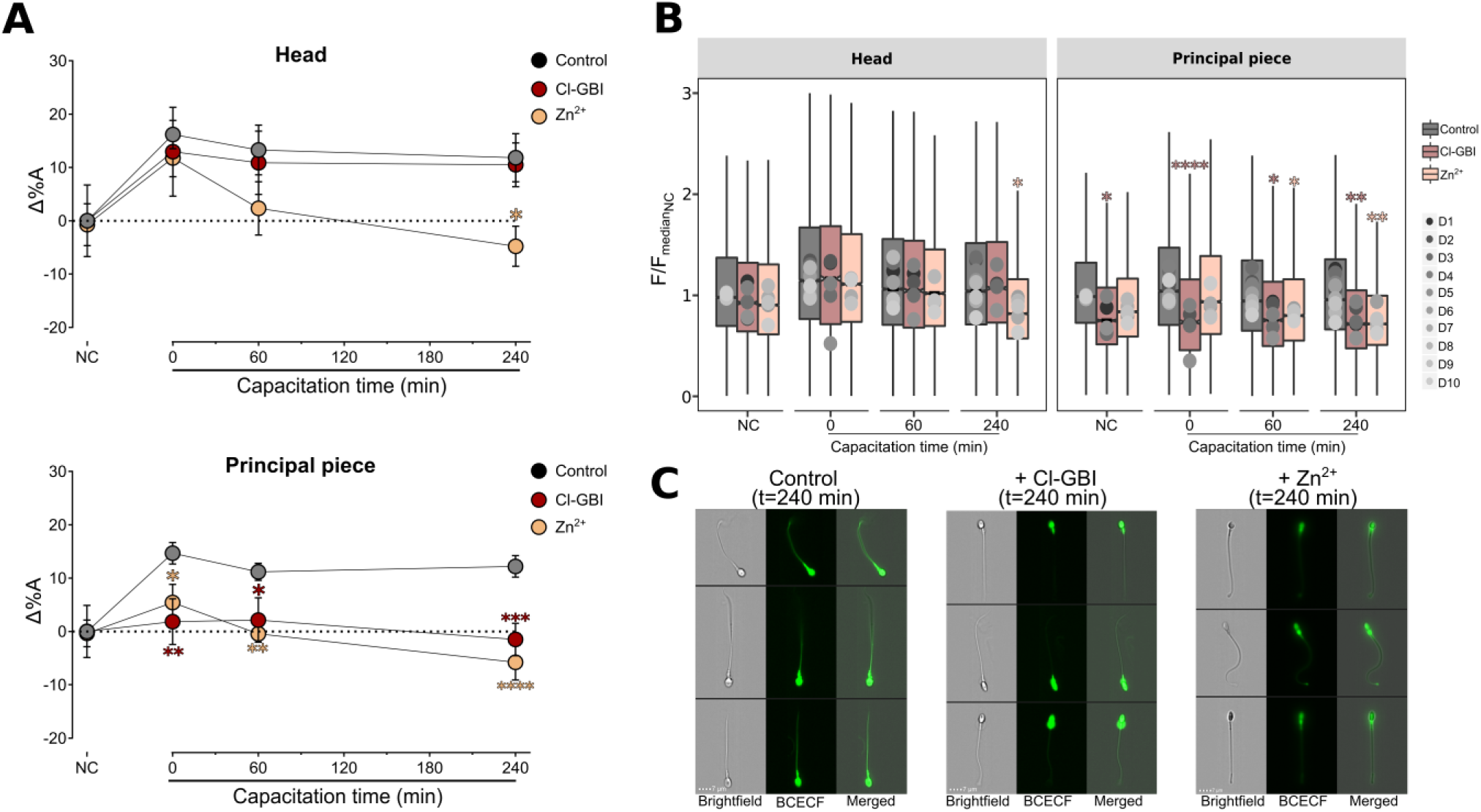
Inhibition of Hv1 prevented alkalization in the principal piece but not in the head. **A.** Δ%A values (Δ%A=%T-%NC_A_) for each subcellular region from human sperm cells placed in non-capacitating medium or in complete capacitating medium and in both media in the presence or absence of 200 µM Cl-GBI or Zn^2+^. Values were measured at the indicated capacitation times. Data represent the mean ± s.e.m. (n=10 control, n=5 for Cl-GBI and Zn^2+^). **B.** Boxplots of normalized BCECF fluorescence values for each subcellular region in cells under the same conditions as in **A**. Gray-scale dots (D1-D10) within boxplots indicate the median fluorescence value for the entire sperm population from each donor, consisting of at least 1,000 cells each (n=10 control, n=5 for Cl-GBI and Zn^2+^). **C.** Three representative images of cells capacitated for 240 min under the indicated conditions. Data were compared using two-way ANOVA with incubation time as one factor, and treatment as the other factor. \**p*<0.05, \*\**p*<0.01, \*\*\**p*<0.001, \*\*\*\**p*<0.0001.

To further confirm Hv1 participation in the pH_i_ increase, we used Zn^2+^, a well-known inhibitor of Hv1. The presence of 200 µM of ZnCl (Zn^2+^) reduced significantly Δ%A in the head at 240 min (*p*=0.0286) of capacitation (Fig. 5A) which was also observed when pooled fluorescence values were analyzed (*p*=0.0245) (Fig. 5B). The pH_i_ increase was strongly inhibited by Zn^2+^ in the principal piece at all capacitation times (*p*<0.0339), the reduction was significant regardless of the analysis (Fig. 5A-B). These data confirm the participation of Hv1 in human sperm pH_i_ regulation, but also corroborate that at least in the head other proteins must be participating in the control of pH_i_ in that cell region.

### Proteins that regulate pH_i_ are required for hyperactivation

The downstream role of HCO_3_^-^ uptake in the control of sperm hyperactivation has been widely recognized (Okamura et al., 1985), primarily via a PKA signaling pathway and CATSPER activation (Orta et al., 2018; Qi et al., 2007; Wennemuth et al., 2003). In this work we showed that Hv1, HCO_3_^-^ influx, and to a lesser extent CFTR, are required for the pH_i_ increases during capacitation. Employing a CASA system, we explored whether inhibition of these proteins and the lack of HCO_3_^-^ affected sperm hyperactivation. Interestingly, none of these conditions produced a change in total motility compared to control conditions during the explored time window (Fig. 6A). In contrast, all these treatments caused, to varying degrees, reduction in the percentage of cells that displayed hyperactivated motility, compared to control conditions. For instance, both Hv1 inhibition and the absence of HCO_3_^-^ in the medium completely prevented hyperactivation (*p*<0.0310) (Fig. 6B). CFTR blocking significantly reduced hyperactivation at 0 min, 60 and 240 min (*p*=0.0193) (Fig. 6B) and inhibition of NBC reduced the number of hyperactived cells (*p*< 0.0001) upon capacitation induction (0 min), but not after 60 and 240 min.

**Figure 6.**
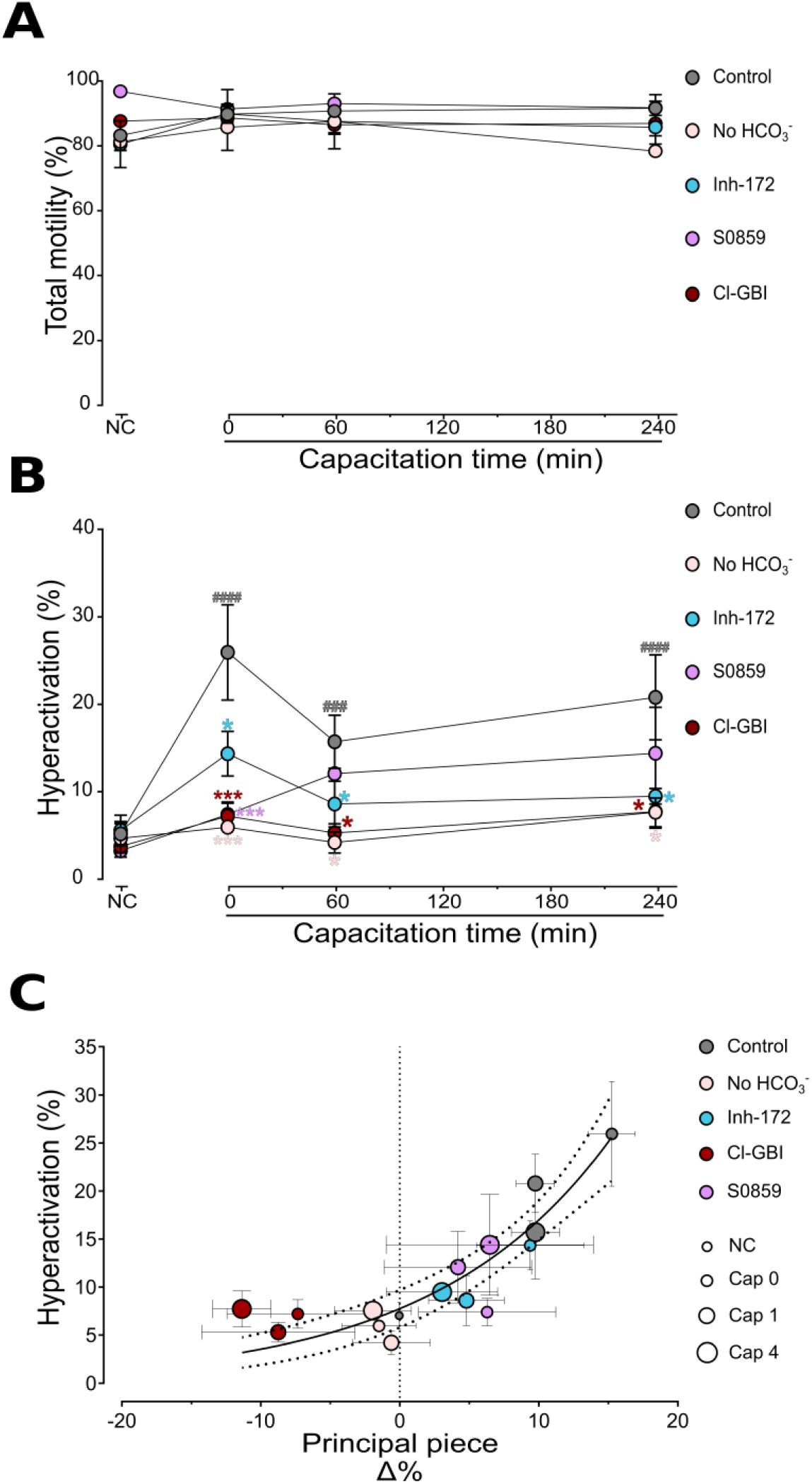
Proteins involved in pH_i_ control are required for hyperactivated motility but not for total motility. **A.** Total motility measurements of human sperm cells placed in non-capacitating medium or in complete capacitating medium and in both media in the presence or absence of 5 µM S0859 or Inh172, or 200 µM Cl-GBI at different times of incubation. Data represent the mean ± s.e.m. (n = 5 for Control and Inh-172, n = 4 for Cl-GBI, S0859 and No HCO_3_^-^). **B.** Quantification of proportion of hyperactive cells during the time and conditions mentioned in A. Data represent the mean ± s.e.m. (n = 5 for Control and Inh-172, n = 4 for Cl-GBI, S0859 and No HCO3 Data were compared using two-way ANOVA with incubation time as one factor, and treatments as the other factor. \**p*<0.05, \*\*\**p*<0.001. ^###^*p*<0.001, ^####^*p*<0.0001. C. Changes in hyperactivation as a function Δ%. The incubation time (NC, capacitation at 0, 60 and 240 min) are depicted as increasing size circles. Solid line indicates the exponential fit (fit parameters are indicated in main text). Dotted line are the 95% confidence intervals. Data represent the mean ± s.e.m.

Lastly, we plotted the percentage of cells exhibiting hyperactivated motility as a function of the percentage of cells with high pp-pH_i_. We found that hyperactivation increases exponentially as a function of alkalization in the principal piece (R^2^=0.80, y = 7.686*e*^(0.007X)^, τ = 12.85). (Fig. 6C).

## DISCUSSION

Through comparisons of initial and final pH_i_, *in vitro* studies have shown that human sperm cells exhibit alkalization after 24 (Cross and Razy-Faulkner, 1997) and 13 (Lopez-Gonzalez et al., 2014) hours of capacitation. More recently, while we were preparing the present manuscript for publication, Brukman and colleagues reported that pH_i_ in a human sperm subpopulation increased slightly after 10 min of capacitation, had a further increase after 1 hour, and then remained constant after 2, 4 and 6 hours (Brukman et al., 2019). These observations were made employing conventional flow cytometry on BCECF-stained cells. For the present study, we applied our recently developed sperm segmentation process using image-based flow cytometry (Matamoros-Volante et al., 2018) to follow human sperm pH_i_ changes in three distinct subcellular regions (head, midpiece and principal piece) at various time points of capacitation (up to 4 hours). We started out by demonstrating that this method can be reliably used to detect pH_i_ changes in all three regions.

As expected, histograms constructed on BCECF fluorescence values measured for each sperm population sample vary in distribution and amplitude across donors and experimental replicates, even under equal treatment conditions. The fluorescence data sets from all donors/replicates were first pooled and displayed as boxplots for every given condition, enabling comparisons and statistical analyses. For each of them, in addition, the percentage of subcellular regions having fluorescence values above those of the third quartile of the NC control was calculated. These values were then plotted as a percentage difference with respect to control conditions in order to display and compare the pH_i_ kinetics during capacitation.

We observed that under conditions that do not support capacitation, pH_i_ in all three subcellular regions remained constant over the 4-h time window. But when sperm were incubated under conditions that promote capacitation, a pH_i_ increase occurred in the head and principal piece, remaining stable during the entire time window studied. Given that no change in midpiece pH_i_ was observed, further analyses of the proteins involved in regulating pH_i_ were conducted solely in the head and principal piece.

Previous evidence obtained by our group suggests that an electrogenic NBC is responsible for HCO_3_^-^ influx during capacitation (Demarco et al., 2003; Puga Molina et al., 2018). In the present work, we observed that pharmacological inhibition of NBC caused a decrease in basal pH_i_ under NC conditions both in the head and principal piece. However, upon addition of capacitation medium, Δ%A was very similar to Δ% in both subcellular regions. These results indicate that NBC participates in pH_i_ homeostasis prior to capacitation, rather than having a role in the pH_i_ increase observed during capacitation, and they also suggest that other proteins are responsible for such increase.

Recently, our group also proposed that HCO_3_^-^ influx might either take place directly through CFTR, or through other transporters coupled to CFTR (Puga Molina et al., 2017; Puga Molina et al., 2018). In the earlier manuscript, based on conventional flow cytometry measurements, we reported that CFTR inhibition causes a decrease in pH_i_ after 5 hours of capacitation. While our present results indicate that CFTR inhibition caused only a slight pH_i_ decrease in the head and principal piece, it was statistically significant in the head at 4 hours of capacitation, in agreement with our previous measurements performed after 5 hours of capacitation. We had previously proposed that the increase in pH_i_ during capacitation could be due to the concerted action of NBC and CFTR (Puga Molina et al., 2017; Puga Molina et al., 2018). Our present results indicate that, at least for the initial pH_i_ increase, NBC and CFTR are not required.

On the other hand, both the absence of HCO_3_^-^ and the general blocking of HCO_3_^-^ transporters completely prevented the pH_i_ increase in both subcellular regions. These results suggest the participation of additional proteins with HCO_3_^-^ transport activity. In this regard, HCO_3_^-^ transporters from the NBC family, such as the electroneutral Na^+^-driven Cl^-^/HCO_3_^-^ exchangers, NDCBE (SLC4A8) and NBCn2 (SLC4A10) have been detected in human testis, albeit only at the transcriptional level (Damkier et al., 2007; Pushkin et al., 2000). Additionally, other proteins related to HCO_3_^-^ transport have been found in mammalian sperm, such as the SLC26 family members A3 and A6 (Chávez et al., 2012; El Khouri et al., 2018), as well as A8 (Touré et al., 2007), and carbonic anhydrase activity has also been detected in human sperm (José et al., 2015; Wandernoth et al., 2010). Further research is needed to investigate whether these other transporters/enzymes are involved in pH_i_ regulation.

The evidence provided here suggests that different proteins are involved in pH_i_ regulation in different sperm subcellular regions. We propose that HCO_3_^-^ transporters in the head (yet to be identified) are responsible for the initial and sustained HCO_3_^-^ uptake. It is then possible that diffusion of HCO_3_^-^ (or a second messenger) occurs from the head to the flagellum, which would explain the delay in pH_i_ increase observed in the principal piece between, compared to the head.

The initial HCO_3_^-^ influx, known to activate a PKA pathway, could presumably also participate in the initial pH_i_ increase. According to previous studies conducted by our group, PKA blockage with H89 causes strong cytoplasmic acidification in capacitated human sperm (Puga Molina et al., 2017). Using this same inhibitor and KH7, our present results corroborate participation of the PKA pathway on pH_i_ regulation, with a major contribution in the principal piece and, to a lesser extent, in the head. PKA localization in human sperm is not restricted to a specific site (Neuhaus et al., 2006), but the main subcellular localization of PKA substrates are in the principal piece (Battistone et al., 2013).

Previous work has established that in human sperm, Hv1 mediates outward H^+^ currents, which are enhanced once sperm are capacitated (Lishko et al., 2010). We found that pharmacological inhibition of Hv1 with both Cl-GBI and Zn^2+^ does prevent alkalization, in the principal piece, but leaves alkalization in the head unaltered. These results are consistent with the reported localization of this channel exclusively in the principal piece (Lishko et al., 2010; Miller et al., 2018). The fact that Hv1 blockage does not prevent the pH_i_ increase in the head, suggests that Hv1 is not the sole pH_i_ regulator in human sperm, and other mechanisms are likely at work in order to generate such alkalization in the head. Unexpectedly, the presence of Zn^2+^ caused a reduction of pH_i_ in the head after 4 hours of capacitation. This effect is presumably due to a Zn^2+^ target other than Hv1, since it was not observed with the specific Hv1 inhibitor Cl-GBI. Although Zn^2+^ is important for sperm physiology, little is known about the Zn^2+^ transporters that operate in human sperm. Nonetheless, the presence of at least some members from the Zip and ZnT families has been described (Foresta et al., 2014). Transport of Zn^2+^ in and out the cell is generally coupled to the transport of another ion. The Zip protein family consists of symporters that couple Zn^2+^ entry together with HCO_3_^-^. Given that the medium used to induce capacitation *in vitro* contains a high concentration of HCO_3_^-^, it is conceivable that the addition of Zn^2+^ enables such cotransport activity to take place. If this is the case, once Zn^2+^ accumulates in the cell, it could potentially be extruded via ZnT transporters. These function as antiporters with H^+^, thereby explaining the observed acidification in the head at 4 hours of capacitation. Interestingly, we observed that inhibition of Hv1 causes a decrease of pH_i_ in the principal piece, even under conditions that do not promote capacitation, suggesting that this channel is also active and participates in pH_i_ regulation prior to capacitation.

Inhibition of Hv1 abolishes alkalization in the principal piece throughout the entire capacitation time window explored, rather than just initially. The existence of a mechanism maintaining Hv1 activity is therefore expected. It has been reported that Hv1 function is upregulated by phosphorylation of some of its serine and threonine residues, presumably by PKC (Hondares et al., 2014; Morgan et al., 2007; Musset et al., 2010). Thus, PKC could potentially be the key player necessary to sustain Hv1 activity during capacitation. PKC is present in human sperm flagella (Kalina et al., 1995), and its activity has been related to sperm motility (Rotem et al., 1990). Additionally, different lines of evidence along with recent work by Brukman and colleagues (Brukman et al., 2019) suggest a possible link between PKA and Hv1 activation during capacitation. Such activation likely involves the participation of other kinases, since direct Hv1 phosphorylation by PKA has not been demonstrated. During capacitation, there is an increase in tyrosine phosphorylation (PY) of different proteins, which occurs downstream of PKA activation, mainly in the sperm tail (Battistone et al., 2013). This process involves the action of at least two different tyrosine kinases (TKs), PYK2 and FER(T) (Alvau et al., 2016; Battistone et al., 2014; Matamoros-Volante et al., 2018). Brukman et al., 2019 showed that the pharmacological inhibition of these TKs blocks the capacitation-associated alkalization in human sperm cells, and they proposed a possible a connection between TKs and PKC, which in turn upregulates Hv1 (Brukman et al., 2019). Evidence from other cell types suggests that H^+^ conductance driven by Hv1 is also affected by PY, for example in granulocytes (Bianchini et al., 1994) and in neutrophils (Nanda and Grinstein, 1995), although the identity of the implicated TKs remains unknown. Altogether, these findings suggest that Hv1 might be regulated by a signaling network involving PKA, PKC and TKs. Additional experiments are needed to support this proposal.

One of the most important downstream effects of HCO_3_^-^ uptake is the induction of a change in sperm motility patterns (Hereng et al., 2014; Wennemuth, 2003). In fact, sperm from infertile patients present low HCO_3_^-^ levels in seminal plasma, which correlates with poor sperm motility (Okamura et al., 1986). HCO_3_^-^ effects on motility are controlled in a Ca^2+^-dependent manner (Ho et al., 2002; Marquez and Suarez, 2007). The sperm-specific alkalization-dependent calcium channel, CATSPER, is the main molecular entity responsible for intracellular [Ca^2+^] changes upon capacitation (Kirichok *et al*., 2006). Genetic ablation of *CatSper* produces infertility because sperm fail to hyperactivate (Qi et al., 2007). In some models, the intracellular alkalization mediated by Hv1 has been proposed to act as a signal that opens CATSPER, in turn triggering and maintaining hyperactivated motility (Lishko and Kirichok, 2010). Besides, proteins of the glycolytic machinery related to ATP production are required to sustain hyperactivation and the dynein-ATPase necessary to axonemal functionality is also pH_i_ dependent (Mannowetz et al., 2012; Ui, 1966). Altogether, the available evidence suggests a tight relationship between pH_i_ and sperm hyperactivation. In the present work, we demonstrate for the first time that pharmacological inhibition of Hv1 reduces hyperactivation, while leaving total motility unchanged. In fact, with the exception of NBC inhibition, the effect on hyperactivation brought about by our experimental treatments always mirrors their effect on pH_i_ in the principal piece. In other words, conditions that completely prevent alkalization in the principal piece (i.e. either Cl-GBI or medium lacking HCO_3_^-^) also reduce hyperactivated motility. Conversely, CFTR inhibition, which elicits a minor decrease on pH_i_, reduces hyperactivation only slightly. Such a correlation is not apparent upon NBC inhibition, as hyperactivation does not occur, even though the pH_i_ increase in the principal piece is similar in magnitude as the one observed under control conditions. In this case, however, mere preincubation with the inhibitor causes an initial reduction in pH_i_ prior to capacitation, and it is thus conceivable that despite alkalization occurring during capacitation, pH_i_ does not reach the necessary threshold to promote hyperactivation. Thus, while we found the relationship between hyperactivation (%) and pH_i_ increase in the principal piece to be exponential, alkalization might need to be high enough to reach a certain threshold in order for hyperactivation to occur.

In summary, we have shown that cytoplasmic [H^+^] in human sperm is differentially controlled in the head and principal regions; this process involves the participation of various proteins, acting under distinct spatiotemporal control mechanisms. Additionally, our results further support the notion that intracellular alkalization plays a key role in the control of sperm motility. The findings reported here highlight the complexity and relevance of pH_i_ dynamics during human sperm capacitation.

## MATERIALS AND METHODS

### Materials

Potassium dihydrogen phosphate (KH_2_PO_4_) and anhydrous glucose were obtained from J.T. Baker (USA). Bovine Serum Albumin (BSA) was purchased from US Biological (USA). 2’,7’-Bis-(2-carboxyethyl)-5-(and-6)-carboxyfluorescein, acetoxymethyl ester (BCECF-AM), MitoTracker Green FM, and propidium iodide (PI) were obtained from Invitrogen (USA). CFTR-Inh-172 was purchased from Calbiochem Inc. (USA). 2-Chloro-N-[[2′-[(cyanoamino) sulfonyl] [1,1′-biphenyl]-4-yl] methyl]-N-[(4-methylphenyl) methyl]-benzamide, known as S0859, was obtained from Cayman Chemical (USA). 2-guanidinebenzimidazole (2-GBI), 5-chloro-2-guanidinebenzimidazole (Cl-GBI), 4,4′-Diisothiocyanatostilbene-2,2′-disulfonic acid disodium salt hydrate (DIDS), N-[2-(p-Bromocinnamylamino)ethyl]-5-isoquinolinesulfonamide dihydrochloride (H89), zinc chloride (Zn^2+^) and (E)-2-(1H-benzo[d]imidazol-2-ylthio)-N′-(5-bromo-2-hydroxybenzylidene) propanehydrazide (KH7) were obtained from Sigma-Aldrich (USA), along with all other chemicals.

### Ethical Approval

Protocols for human sperm use were approved by the Bioethics Committee of the Instituto de Biotecnología (UNAM, México). Informed consent forms were signed by all donors.

### Culture media

The non-capacitating (NC) medium used in this study was HEPES-buffered Human Tubal Fluid (HTF) Solution containing (in mM): NaCl 90.69, KCl 4.67, CaCl_2_ 1.6, MgSO_4_ 1.2, KH_2_PO_4_ 0.314, Glucose 2.78, Na-Pyruvate 3.38, Na-lactate 60, Hepes 23.8. Capacitation-inducing conditions consisted of HTF medium supplemented with 25 mM NaHCO_3_ and 5% BSA (w/v). All media were adjusted to pH 7.4 with HCl, and the osmolarity was maintained at around 290 mOs kg^-1^.

### Sperm

Sperm samples were obtained from healthy donors, collected by masturbation after 3-5 days of sexual abstinence. Only those samples with normal seminal parameters (according to the 2010 WHO criteria) were used in the study. Semen samples were liquefied for 30 min at 37°C under an atmosphere of 5% CO_2_ in air. Motile sperm were obtained by the swim-up technique, employing NC medium for 1 h at 37°C under an atmosphere of 5% CO_2_ in air. A Makler® Counting Chamber (Sefi Medical Instruments, Israel) was used to adjust the sperm concentration at 10×10^6^ cells/mL. Sperm samples in NC medium were loaded with 250 nm BCECF-AM (see below) and then incubated during 15 min at 37°C, protected from light. Excess dye was removed by centrifugation at 300 g for 5 min, and the cell pellet was resuspended in NC medium to obtain a sperm concentration of 2-8 × 10^6^ cells/mL. For pharmacology evaluations, these BCECF-AM-loaded cells were pre-incubated for 10 min with the various blockers tested (5 µM of either Inh-172 or S0859, 100 µM DIDS, 50 µM of KH7, 30 µM H89, 200 µM of Cl-GBI and 200 µM Zn^2+^) or with the vehicle (DMSO or culture media) alone (control). After pre-incubation, an aliquot of cells was combined with an equal volume of 2X capacitation medium (supplemented with the same concentrations of the different blockers), and sperm cells were incubated at 37°C under a 5% CO_2_ atmosphere. At different time periods, up to a maximum capacitation time of four hours, an aliquot of cells was taken for analysis.

### Intracellular pH estimation by image-based flow cytometry

Intracellular pH (pH_i_) was monitored through fluorescence measurements using the pH-sensitive cell-permeable probe BCECF-AM. Once this dye enters the cell, cytosolic estereases cleave the acetoxymethyl ester (AM) group and free BCECF accumulates in the cell’s cytosol. The intensity of this dye’s fluorescence emission (λ=535 nm) increases with increasing pH, enabling the tracking of pH_i_ conditions. BCECF-loaded cells, in either NC or capacitation medium were concentrated from 2 × 10^6^ to 8 × 10^6^ cells/mL (in a final volume of 50 µL) by centrifugation at 300 g for 5 min. At least 30 seconds before measurements, 250 nM PI (final concentration) was added to the cell suspension to evaluate sperm viability. BCECF and PI fluorescence were measured using the image-based flow cytometer ImageStream Mark II (Amnis, USA). The acquisition settings of INSPIRE® software (Amnis, USA) were as follows: objective: 60X magnification, excitation laser: 488nm, laser intensity range: 20-100 mW (in order to avoid over excitation and pixel saturation), BCECF emission: 535nm, collected in channel 2 (range 480-560 nm), PI emission: 620 nm, collected in channel 4 (range 595-660 nm), brightfield images: channel 1. During acquisition, different parameters were set for preliminary discrimination of saturated, cell aggregates, non-sperm (*e.g*. round cells), and non-focused cells according to previous work (Matamoros-Volante et al., 2018). After pre-processing, 12,500 cells were recorded for each condition.

To estimate the kinetics of pH_i_ changes during capacitation, we employed the aforementioned conditions to assess BCECF fluorescence under NC conditions (*i.e.* after swim-up and dye loading). For measurements under capacitation conditions, recordings were done immediately after the addition of 2X capacitation medium (considered capacitation time = 0 min). Thereafter, we tracked pH_i_ under capacitation conditions at 15-min intervals, up to 180 min of incubation, unless otherwise specified. A final measurement was made at 240 min of capacitation.

### Computer assisted sperm analysis (CASA)

The effect of the various blockers in sperm motility was evaluated using a CASA system. A 7-µL aliquot of each sperm sample was placed in a pre-warmed microscopy slide, covered with a coverslip (18 × 18 mm), and sperm motility was monitored using a negative phase contrast 10X objective (Nikon, Japan). Data was acquired using Sperm Class Analyzer software (SCA, Microptics, Barcelona, Spain). 500 cells were measured for each experimental condition, by collecting 25 images with a frequency of 50 Hz. Sperm hyperactivation was assessed according to the criteria established by Mortimer, 2000, as follows: curvilinear velocity (VCL): > 150 µm/s; linearity (LIN): < 50%; half lateral head displacement (ALH_1/2_): > 3.5 µm.

### Image based-flow cytometry data analysis

Image-based flow cytometry-derived images were analyzed with IDEAS® software version 6.2 (Amnis, USA) using a previously reported analysis strategy designed to: a) discriminate non-sperm events (doublets, debris, etc.), unfocused images, and dead cells (positive to PI); and b) perform segmentation of sperm images in order to selectively analyze three distinct sperm cell regions, namely the head, the midpiece, and the principal piece (Matamoros-Volante et al., 2018). After completing the formerly described selection and segmentation processes, anywhere between 1,000 and 2,000 cell regions per treatment remained for analysis from each of the semen samples (n=3 to 9). For each treatment, fluorescence histogram data (*e.g*. Fig.S1A) from the various semen samples were pooled into boxplots (*e.g*. Fig. S1B) for each of the three analyzed cell regions. In order to identify pH_i_ increases across sperm populations subjected to various treatments and/or capacitation time points, only cell regions exhibiting a fluorescence value higher than those of the third quartile in the NC condition (i.e. falling to the right of the dashed line in Fig. S1A-B) were arbitrarily considered as having a high pH_i_. The percentage of such high pH_i_ cell regions (Fig. S1B) was then calculated for the NC condition (%NC) and for each treatment/time point (%T), after eliminating outliers (i.e. those with fluorescence values in the top 5%). To assess the effect of each capacitation time point (%T) or treatment (%T_A_), the difference between those percentages (Δ%) was calculated as follows: Δ%=%T-%NC (*e.g*. Fig. S1C). In many cases, the %NC value under altered conditions (%NC_A_) (i.e. absence of HCO_3_^-^ or presence of a blocker) was significantly lower than that of %NC (*e.g*. Fig. S1B-C), resulting in negative Δ% values (*e.g.* Fig. S1C). To enable side-by-side comparisons of pH_i_ kinetics, we adjusted Δ% values for altered conditions to start at zero (*i.e.* equal to the NC condition) through an alternative calculation: Δ%A =%T_A_ - %NC_A_ (*e.g.* Fig. S1D). With this normalization, the effect of each treatment is measured with respect to its corresponding initial NC condition (NC or NC_A_). However, in order to appreciate the effect of pre-incubation with blockers on the initial %NC_A_ value, we also show all pHi kinetics plots using %NC for all Δ% calculations (Fig. S2A-D, see corresponding normalized results in Figures 2A, 3A, 4A and 5A).

### Statistical analysis

Results from image-based flow cytometry are presented as boxplots of the pooled fluorescence values for all analyzed cell regions from all donors using the median fluorescence value at each condition divided by the median fluorescence value of the NC condition (*e.g*. Fig S1B). The corresponding calculated Δ% and Δ%A values are presented as mean +/- s.e.m. Differences in these values were assessed using two-way ANOVA, considering capacitation time (*e.g*. NC, 0, 60, 240 min, etc.) as one factor, and treatment (*e.g.* Control, Inh-172, Cl-GBI, etc.) as the second factor. Motility measurements are presented as mean +/- s.e.m. and statistical differences assessed also with two-way ANOVA. The Tukey test was subsequently applied to determine differences between treatments. A probability (*p*) value <0.05 was considered a statistically significant difference. GraphPad Prism version 6 (GraphPad, USA) was used for statistical analysis. ggplot2 library (Wickham, 2009) in R studio software (R Core Team, 2017) was employed for plotting and data analysis. The final versions of the figures were prepared using Inkscape 0.91 (Inkscape.org, USA).

## ACKNOWLEDGMENTS

We thank Paulina Torres, Andres Saralegui, Yoloxóchilt Sánchez and Jose Luis De la Vega for technical assistance. We thank Shirley Ainsworth for library support. We acknowledge Juan Manuel Hurtado, Roberto Rodríguez, Omar Arriaga and Arturo Ocádiz for computer services. We thank Marcela Trevino for critically reading the manuscript and English editing.

